# The Ebola virus interferon antagonist VP24 undergoes active nucleocytoplasmic trafficking

**DOI:** 10.1101/2020.08.10.245563

**Authors:** Angela R. Harrison, Gregory W. Moseley

## Abstract

Viral interferon (IFN) antagonist proteins mediate evasion of IFN-mediated innate immunity and are often multifunctional, having distinct roles in viral replication processes. Functions of the Ebola virus (EBOV) IFN antagonist VP24 include nucleocapsid assembly during cytoplasmic replication and inhibition of IFN-activated signalling by STAT1. For the latter, VP24 prevents STAT1 nuclear import *via* competitive binding to nuclear import receptors (karyopherins). Many viral proteins, including proteins from viruses with cytoplasmic replication cycles, interact with the trafficking machinery to undergo nucleocytoplasmic transport, with key roles in pathogenesis. Despite established karyopherin interaction, the nuclear trafficking profile of VP24 has not been investigated. We find that VP24 becomes strongly nuclear following overexpression of karyopherin or inhibition of nuclear export pathways. Molecular mapping indicates that cytoplasmic localisation of VP24 depends on a CRM1-dependent nuclear export sequence at the VP24 C-terminus. Nuclear export is not required for STAT1 antagonism, consistent with competitive karyopherin binding being the principal antagonistic mechanism while export mediates return of nuclear VP24 to the cytoplasm for replication functions. Thus, nuclear export of VP24 might provide novel targets for antiviral approaches.

**Importance:** Ebola virus (EBOV) is the causative agent of ongoing outbreaks of severe haemorrhagic fever with case-fatality rates between 40 and 60%. Proteins of many viruses with cytoplasmic replication cycles similar to EBOV interact with the nuclear trafficking machinery, resulting in active nucleocytoplasmic shuttling important to immune evasion and other intranuclear functions. However, exploitation of host trafficking machinery for nucleocytoplasmic transport by EBOV has not been directly examined. We find that the EBOV protein VP24 is actively trafficked between the nucleus and cytoplasm, and identify the specific pathways and sequences involved. The data indicate that nucleocytoplasmic trafficking is important for the multifunctional nature of VP24, which has critical roles in immune evasion and viral replication, identifying a new mechanism in infection by this highly lethal pathogen, and potential target for antivirals.

## Introduction

*Zaire ebolavirus*, commonly known as Ebola virus (EBOV), is a causative agent of multiple outbreaks of Ebola severe haemorrhagic fever, including the 2014-2016 West African outbreak and global health emergency, and the recent ongoing outbreak in the Democratic Republic of Congo. EBOV and other members of the *Ebolavirus* genus belong to the family *Filoviridae*, which also includes another human pathogen, Marburg virus (MABV; genus *Marburgvirus*), and Lloviu virus (LLOV; genus *Cuevavirus*) that was identified in bats in 2011 (1). Filoviruses belong to the order *Mononegavirales* and so have a non-segmented negative-sense RNA genome; transcription and replication of the genome is exclusively cytoplasmic (2).

VP24 is one of the seven genes typically encoded in the filovirus genome (2) and was originally designated a secondary matrix protein (3). However, accumulating evidence indicates critical roles in genome packaging and the formation, condensation and intracytoplasmic transport of nucleocapsids (4-14). Consistent with this, VP24 localises to cytoplasmic inclusion bodies during infection, which are the sites of replication and nucleocapsid formation (14-16). EBOV VP24 also functions as an interferon (IFN) antagonist to suppress the type I IFN-mediated innate antiviral immune response. Specifically, VP24 blocks the nuclear accumulation of the IFN-activated transcription factor STAT1, the key mediator of IFN signalling, by binding competitively to specific karyopherin nuclear import receptors that are responsible for STAT1 transport (17-20).

Entry to the nucleus is restricted by the impermeable double-membrane nuclear envelope, such that all nucleocytoplasmic transport occurs through nuclear pore complexes (NPCs) embedded in the envelope. Proteins/molecules smaller than c. 40-65 kDa are able to diffuse through the NPC, but specific directional transport of protein cargoes is mediated by expression of nuclear localisation and nuclear export sequences (NLSs and NESs) that bind to members of the karyopherin family (also known as importins or exportins) in the cytoplasm or nucleus. Karyopherins mediate energy-dependent translocation of cargo through the NPC (reviewed in (21)); this enables regulable nucleocytoplasmic localisation of proteins, and is absolutely required for transport of cargoes larger than the diffusion limit. In the cytoplasm, karyopherin alpha (Kα) adaptor proteins typically recognise NLSs comprising mono- or bi-partite sequences enriched in positively-charged residues. Kαs also bind to karyopherin beta (Kβ), which facilitates movement of the cargo/Kα/Kβ complex through the NPC. Kβs can additionally directly bind and facilitate import of cargoes containing certain NLSs. Within the nucleus, cargoes containing NESs (commonly a motif of hydrophobic residues) bind to exportins, including the ubiquitously expressed and well-characterised chromosomal maintenance 1 (CRM1), to be exported to the cytoplasm (21).

STAT1 does not contain a classical NLS and uses a conformational NLS that is presented on IFN-activated parallel STAT1 homodimers and STAT1-STAT2 heterodimers to mediate nuclear import, which enables transcriptional activation of IFN-stimulated genes (22). STAT1 dimers bind to members of the NPI-1 Kα sub-family (Kα1, 5 and 6) at a specific site distinct from sites bound by other cellular cargoes containing classical NLSs (18, 23, 24). VP24 binds competitively to this site, inhibiting nuclear import of STAT1, as well as other cellular cargoes that use the same site, but not cargoes that bind elsewhere (17, 18, 20, 25). The interaction between VP24 and Kα has been the subject of intense research, revealing a well-defined VP24-Kα interface involving contact *via* three clusters of residues in VP24, with importance to IFN antagonism (19, 20) and VP24 stability (26).

Several IFN antagonists use karyopherin binding to inhibit nuclear import of host cargo. Severe acute respiratory syndrome coronavirus (SARS-CoV) ORF6 protein tethers Kβ1/Kα2 complexes to the ER/golgi membrane, inhibiting IFN-induced STAT1 nuclear import (27). 4b protein of Middle East respiratory syndrome coronavirus (MERS-CoV) contains a classical bipartite NLS that competes with nuclear factor κβ (NF-κβ) for Kα4 to inhibit NF-κβ-dependent expression of pro-inflammatory cytokines (28). While SARS-CoV ORF6 remains cytoplasmic due to ER/Golgi association (27), the 4b NLS-Kα4 interaction mediates 4b nuclear import, such that this protein is predominantly nuclear (28). Hijacking of nuclear trafficking pathways for import/export of proteins is common in viruses with cytoplasmic replication cycles, including RNA viruses such as MERS-CoV (28), rabies virus (RABV) (29) and henipaviruses (30, 31), and these processes have been linked to pathogenesis. For example, the RABV IFN antagonist P1 protein binds to STAT1 and shuttles *via* multiple NLSs and NESs, but is largely cytoplasmic due to a dominant NES that effects nuclear exclusion of P1-STAT1 complexes. Defective nuclear export of P1 correlates with impaired IFN antagonism and viral attenuation (32-34). Many proteins of cytoplasmic viruses also form intranuclear interactions/functions, including isoforms of RABV P protein and the matrix proteins of Nipah and Hendra viruses, enabling cytoplasmic viruses to modulate intranuclear processes (35-37). However, despite extensive characterisation of the VP24-Kα interaction, the nuclear trafficking profile of VP24 remains unresolved.

Here, we report that EBOV VP24 can undergo specific trafficking between the nucleus and cytoplasm, involving a C-terminally located NES that enables CRM1-dependent nuclear export. By identifying critical residues in the NES, we find that VP24 nuclear export is not essential for STAT1 antagonist function, consistent with competitive Kα binding as the key mechanism, and so appears to be required due to the multifunctional nature of VP24 that involves cytoplasmic roles in the replication cycle, distinct from immune evasion.

## Results

### EBOV VP24 undergoes nucleocytoplasmic trafficking

EBOV VP24 is largely excluded from the nucleus in infected cells (16, 38) and is cytoplasmic or diffuse in transfected cells (17, 19, 20, 39), such that VP24 differs from MERS-CoV 4b, which accumulates within the nucleus (28), but is similar to SARS-CoV ORF6, which is largely cytoplasmic (27). Since VP24 binds to Kαs at a site overlapping the site that mediates active nuclear import of STAT1 (20), it appears likely that cytoplasmic localisation is due to physical sequestration (similar to ORF6) or rapid nuclear export following entry to the nucleus. We thus examined whether VP24/Kα1 complexes can accumulate within the nucleus by expressing full-length VP24 (residues 1-251) fused to GFP (GFP-VP24) or GFP alone in COS7 cells, with or without co-expression of FLAG-tagged Kα1 or a FLAG control; cells were then immunostained for FLAG and imaged by confocal laser scanning microscopy (CLSM) (Figure 1A). Nucleocytoplasmic localisation of the proteins was quantified by calculating the nuclear to cytoplasmic fluorescence ratio (Fn/c, Figure 1B), as previously (34, 40).

**Figure 1.**
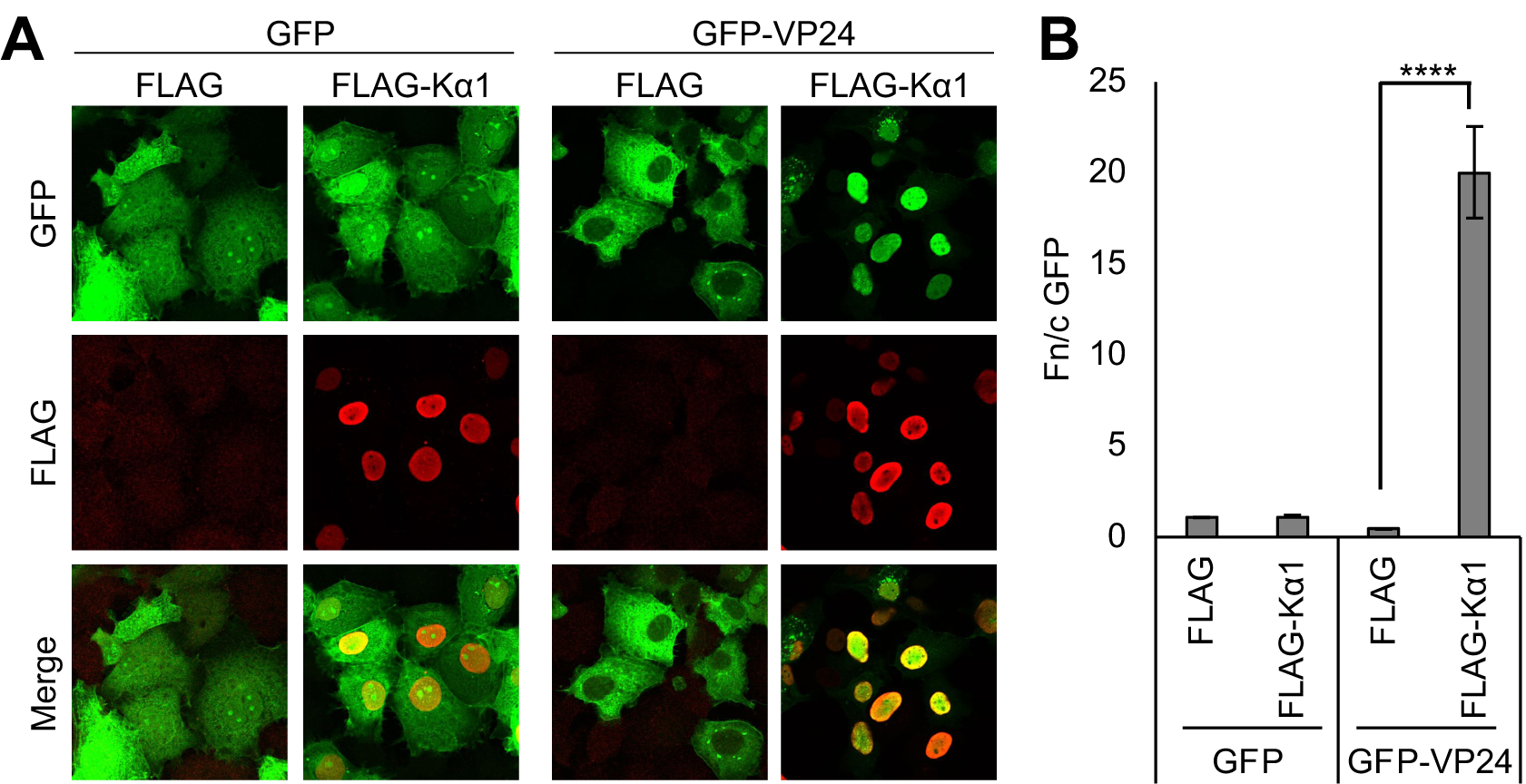
Kα1-VP24 complexes can localise to the nucleus. (A) COS7 cells co-transfected to express GFP or GFP-VP24 with FLAG (control) or FLAG-Kα1 were fixed 24 h post-transfection before immunofluorescent staining for FLAG (red) and CLSM analysis. Representative images are shown. (B) Images such as those shown in A were analysed to calculate the nuclear to cytoplasmic fluorescence ratio (Fn/c) for GFP (mean ± SEM, n ≥ 52 cells for each condition; results are from a single assay representative of three independent assays). Statistical analysis (Student’s *t*-test) was performed using GraphPad Prism software. ****, p < 0.0001.

FLAG-Kα1 localised strongly to the nucleus (as expected (27)), irrespective of VP24 expression, and Kα1 expression had no apparent effect on localisation of GFP alone. GFP-VP24 could be detected in the nucleus and cytoplasm in cells co-expressing the FLAG control, but localised predominantly to the cytoplasm (Figure 1A,B), consistent with studies in infected cells (16, 38) and cells expressing HA- or GFP-tagged VP24 (19, 39). However, Kα1 over-expression effected strong translocation of GFP-VP24 into the nucleus, resulting in clear intranuclear co-localisation of GFP-VP24 and FLAG-Kα1. Consistent with previous reports of Kα1-VP24 interaction (17, 18), FLAG-Kα1 co-precipitated with GFP-VP24 from HEK293T cells (Figure S1). Thus, GFP-VP24 can localise into the nucleus in complexes with Kα1, indicating that cytoplasmic localisation, which is required for roles in nucleocapsid assembly/condensation (4, 5, 7), derives from active nuclear export.

To assess the role of cellular nuclear export pathways in VP24 localisation, we examined the effect on VP24 of leptomycin B (LMB), an inhibitor of the exportin CRM1 (32, 40). COS7 cells expressing GFP or full-length GFP-VP24 (Figure 2A) were treated with or without LMB before imaging live by quantitative CLSM (Figure 2B,C). As expected, GFP (∼ 30 kDa), which can diffuse through the NPC and lacks NLSs or NESs, was diffusely localised between the cytoplasm and nucleus, with negligible effect of LMB. GFP-VP24^1-251^ was predominantly cytoplasmic at steady state in living cells, consistent with localisation of VP24 in fixed cells (Figure 1) (19, 39). Following LMB treatment, GFP-VP24^1-251^ clearly re-localised from the cytoplasm to the nucleus (> 4 fold increase in Fn/c), indicating that VP24 undergoes active export from the nucleus mediated by CRM1.

**Figure 2.**
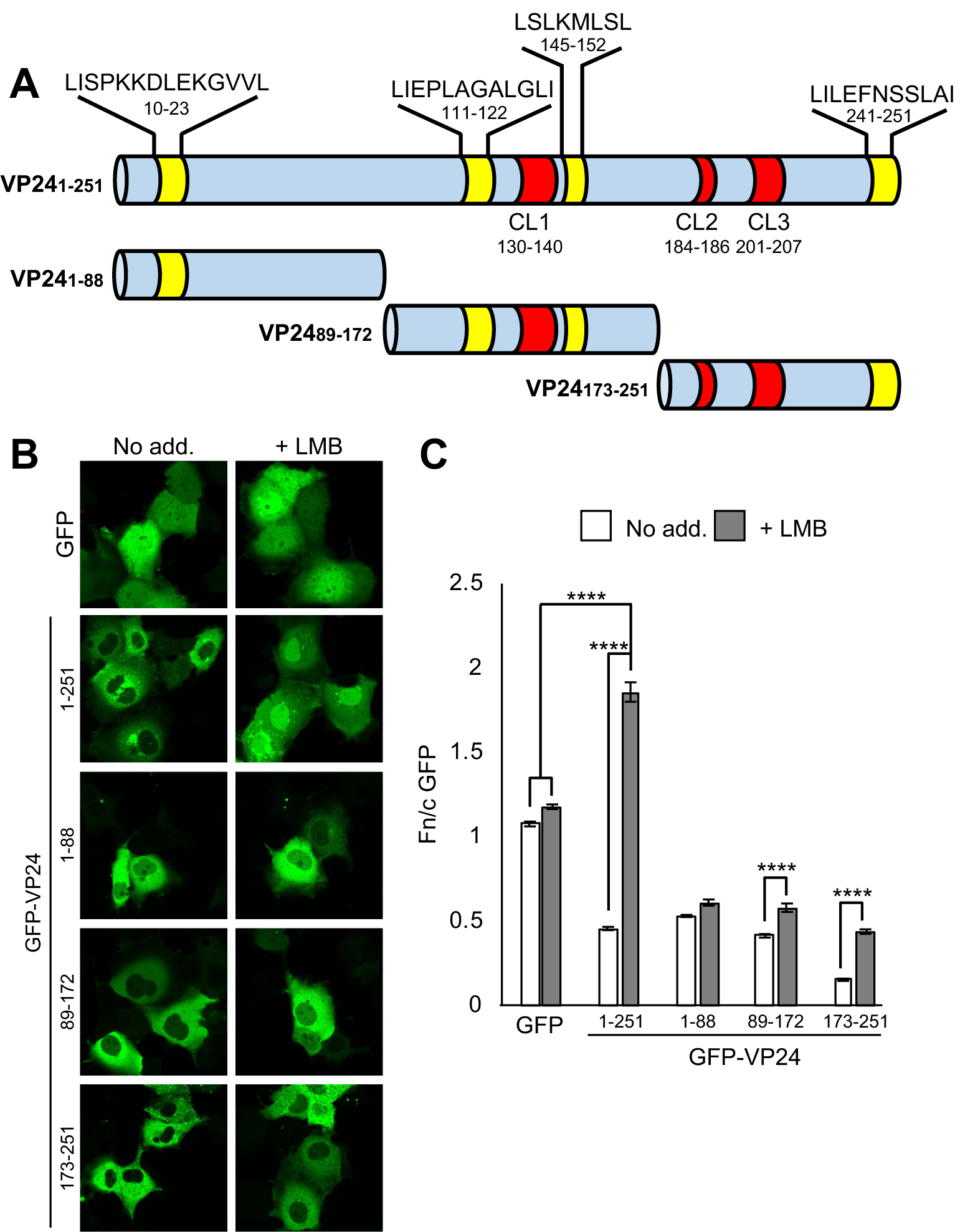
EBOV VP24 undergoes CRM1-dependent nuclear export. (A) Schematic of full-length VP24 and truncated VP24 proteins generated. Location of potential NESs are shown in yellow. Location of clusters (CL1-3) of residues that interact with Kαs in the VP24:Kα5 complex crystal structure (20) are shown in red. Numbering indicates residue positions in full-length VP24; sequences of potential NESs are shown above. (B) COS7 cells transfected to express the indicated proteins were treated 24 h post-transfection with or without LMB (2.8 ng/ml, 3 h) before live-cell CLSM analysis. Representative images are shown. (C) Images such as those shown in B were analysed to calculate the Fn/c for GFP (C; mean ± SEM; n ≥ 31 cells for each condition; results are from a single assay representative of three independent assays). Statistical analysis used Student’s *t*-test. ****, p < 0.0001; No add., no addition.

CRM1 facilitates nuclear export of a broad range of cellular and viral cargoes (including RABV P1, Hendra virus matrix protein, Measles virus C protein (31)) that present NESs typically conforming to a motif of hydrophobic residues (L-X^(2-3)^-L-X^(2-3)^-L-X-L, where L corresponds to L, V, I, F or M, and X is any amino acid (21)). Manual inspection of the VP24 sequence and analysis using the online NES prediction server NetNES (41) identified four potential CRM1-dependent NESs (Figure 2A, Figure S2A). Importantly, the Fn/c for GFP-VP24^1-251^ in LMB-treated cells was higher than that for GFP alone (Figure 2C), indicative of accumulation. Thus, VP24 localisation appears to be dynamic, involving nuclear entry and rapid nuclear export *via* CRM1 interaction.

### EBOV VP24 incorporates a CRM1-dependent NES in the C-terminus

To determine which of the predicted NESs is/are responsible for nuclear export, we generated constructs to express truncated VP24 proteins comprising N-terminal (VP24^1-88^), central (VP24^89-172^) and C-terminal (VP24^173-251^) portions fused to GFP; each of these contained one or more of the potential NESs (Figure 2A). The truncated proteins were designed to be of similar length and to avoid disruption of key structural elements (e.g. alpha helices and beta sheets), based on the VP24 crystal structure (20). All proteins were predominantly cytoplasmic at steady state (Figure 2B). Localisation of the N-terminal fragment was largely unaffected by LMB treatment, and LMB produced only a small (≤ 1.4 fold) increase for the Fn/c of the central fragment (Figure 2B,C). In contrast, a consistent and substantial increase (> 2 fold) in the Fn/c for the C-terminal fragment was observed following LMB treatment. VP24^173-251^ also displayed a consistently reduced Fn/c at steady state compared with the other truncated proteins. Thus, it appeared that prominent discrete CRM1-dependent NES activity is located in the C-terminal region of VP24.

Notably, only full-length VP24 displayed accumulation into the nucleus following LMB treatment, with all truncated proteins remaining significantly less nuclear than GFP alone. This suggests that the full protein sequence is required for efficient nuclear accumulation, such that truncations remove key sequences or otherwise impact conformation to affect important interactions. The crystal structure of VP24 bound to Kα5 indicates that three regions contact the Kα (CL1 and CL2/3, separated by 40-60 residues, Figure 2A), and the importance of these in Kα binding was confirmed by mutagenesis (20). Thus, efficient Kα interaction is likely to be impacted in the truncated proteins, as each lacks at least one CL sequence. Other sequences involved in distinct cytoplasmic or nuclear interactions are also likely to contribute, and might be removed or affected by truncation. Thus, to directly confirm that the indicated NES sequences(s) have classical NES activity in terms of being able to re-localise NLS-containing proteins, we generated constructs in which VP24^89-172^ and VP24^173-251^ are fused to an exogenous classical Kα/Kβ-binding NLS from human cytomegalovirus UL44 protein (42), used previously to confirm NES activity in RABV P protein (43). We also generated VP24^89-251^, which contains all CL sequences and the indicated NES sequences (Figure 2A).

CLSM analysis indicated that fusion of the UL44 NLS to GFP (GFP-UL44^NLS^) results in a modest increase in nuclear accumulation, as expected (Figure 3A,B) (42). Fusion of GFP-UL44^NLS^ to VP24^89-172^ or VP24^173-251^ significantly reduced nuclear localisation, consistent with nuclear export and/or cytoplasmic arrest. Similar to GFP-VP24^89-172^ alone (Figure 2B,C), LMB induced only a small increase in Fn/c for GFP-UL44^NLS^-VP24^89-172^ (Figure 3A,B), suggestive of cytoplasmic retention or nuclear export mediated largely *via* an alternative mechanism to CRM1-dependent export. However, LMB induced substantial nuclear localisation of GFP-UL44^NLS^-VP24^173-251^ (> 4.6 fold increase in Fn/c; Figure 3B) that clearly exceeded nuclear localisation of GFP-VP24^173-251^ (Figure 2C), consistent with a classical CRM1-dependent NES counteracting the activity of the heterologous UL44 NLS. The Fn/c for GFP-VP24^89-251^ was also markedly increased by LMB treatment but did not attain an Fn/c similar to that of full-length GFP-VP24 (Figure 2C, Figure 3B), indicating that the complete protein sequence is required for efficient nuclear localisation. Nevertheless, these data clearly indicate that VP24 contains classical CRM1-dependent NES activity and that the principal NES is within VP24^173-251^.

**Figure 3.**
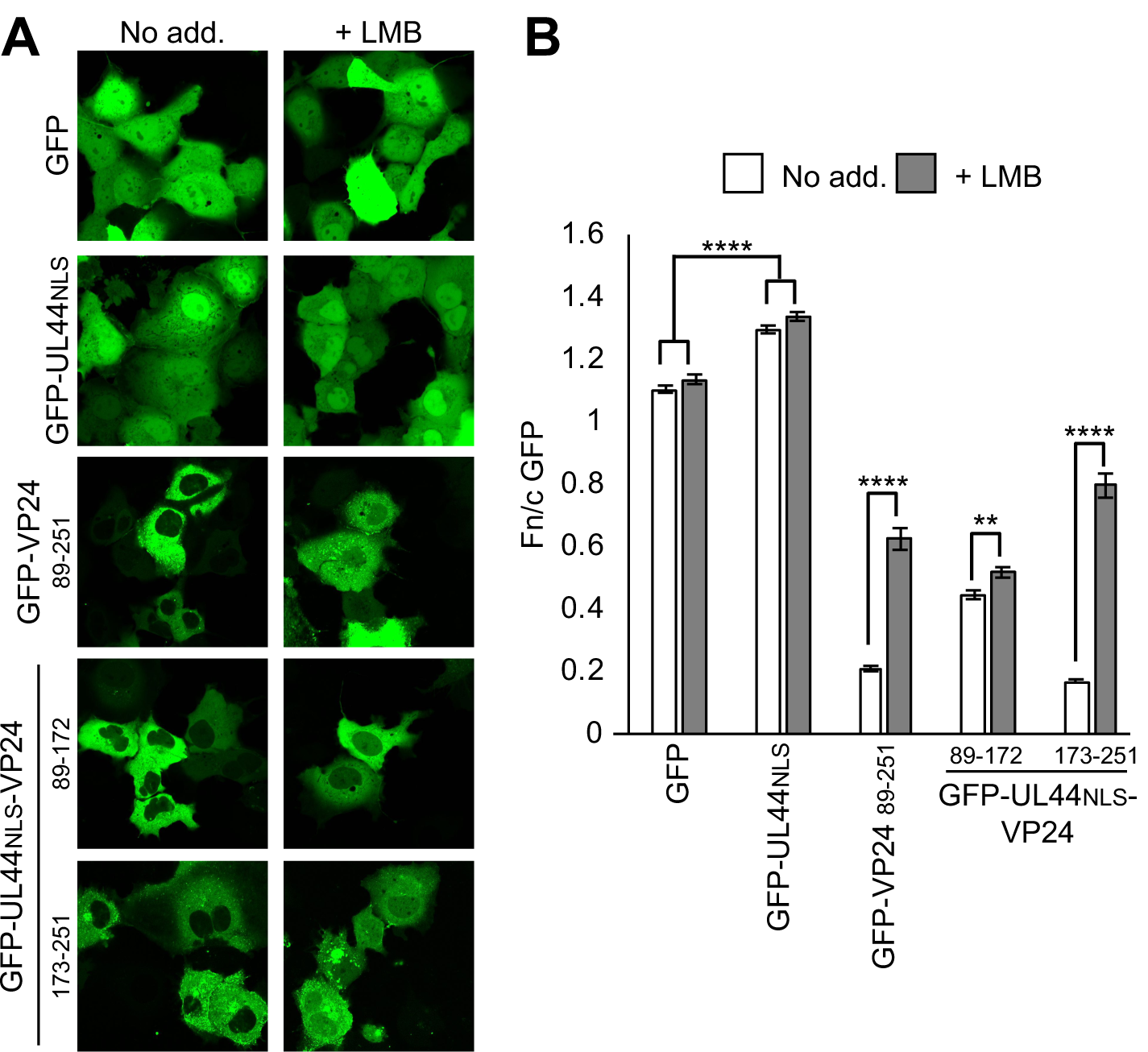
EBOV VP24 C-terminal region contains discrete CRM1-dependent NES activity. (A,B) COS7 cells transfected to express the indicated proteins were treated 24 h post-transfection with or without LMB (2.8 ng/ml, 3 h) before live-cell CLSM analysis (A) and determination of the Fn/c for GFP (B; mean ± SEM; n ≥ 40 cells for each condition; results are from a single assay representative of two independent assays). Statistical analysis used Student’s *t*-test. **, p < 0.01; ****, p < 0.0001; no add., No addition.

### The C-terminal CRM1-dependent NES is the principal sequence mediating nuclear export of VP24

To confirm that the C-terminal NES is the major sequence driving CRM1-dependent export of VP24, we used site-directed mutagenesis to disable the NES motif. Analysis of the VP24 C-terminal region identified residues 241-251 (comprising the C-terminal 11 residues) as containing a sequence strongly conforming to a NES (Figure 2A), with L243, F245 and L249 having the highest NetNES scores among hydrophobic residues in the region (Figure S2A). *In silico* substitution of these residues to alanine (termed NES mutant, NM) abolished the predicted NES (Figure S2B).

Introduction of the substitutions to full-length VP24 significantly enhanced nuclear localisation, resulting in an Fn/c equivalent to that for WT VP24 in LMB-treated cells (Figure 4A,B). Furthermore, the mutations entirely ablated effects of LMB. Thus, L243A/F245A/L249A mutations are sufficient to disable CRM1-dependent nuclear export of VP24, identifying these residues as critical elements of a novel VP24 NES. Similar analysis of GFP-UL44^NLS^-VP24^173-251^ produced comparable results, with the mutations resulting in a significant increase in nuclear accumulation in untreated cells, attaining an Fn/c equivalent to that observed for the WT protein in LMB-treated cells (Figure 4C,D). LMB treatment had only a minor residual effect on the localisation of the mutated GFP-UL44^NLS^-VP24^173-251^ compared with the WT protein, consistent with mutations largely disabling nuclear export activity in the truncated protein.

**Figure 4.**
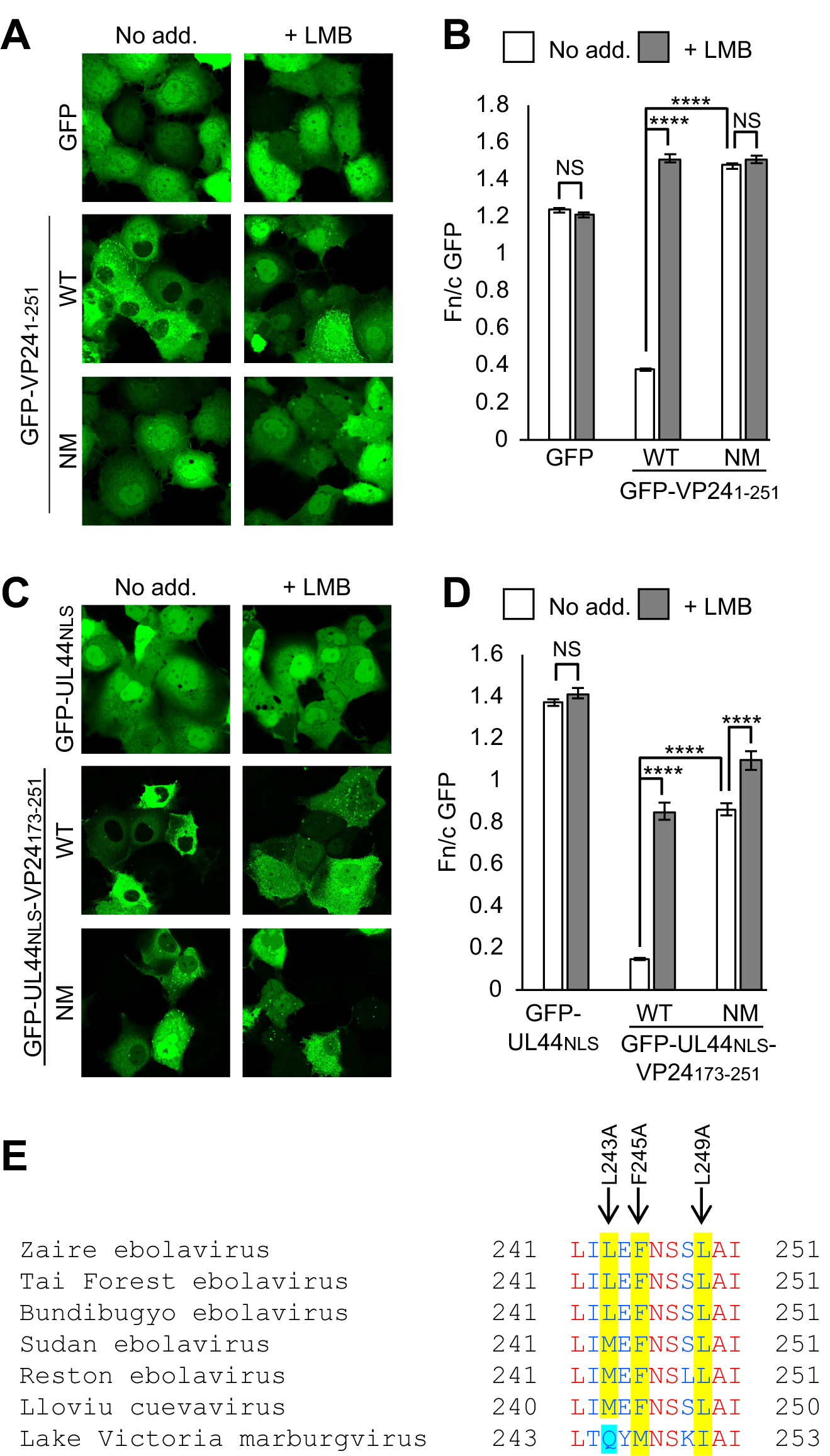
VP24 residues L243, F245 and L249 are critical for VP24 nuclear export. (A-D) COS7 cells transfected to express the indicated proteins were treated 24 h post-transfection with or without LMB (2.8 ng/ml, 3 h) before live-cell CLSM analysis (A,C) and determination of the Fn/c for GFP (B,D; mean ± SEM; n ≥ 50 cells for each condition; results are from single assays representative of three independent assays). Statistical analysis used Student’s *t*-test. ****, p < 0.0001; NS, not significant; No add., no addition; WT, wildtype; NM, NES mutant. (E) Alignment of the C-terminal 11 residues of VP24. Red and blue font indicate conserved and non-conserved residues, respectively. EBOV VP24 residues implicated in CRM1-dependent nuclear export (A-D) are highlighted in yellow. Q245 of MABV VP24, which results in loss of NES consensus, is highlighted in blue.

Comparison of the C-terminal 11 residues of VP24 from species of the *Ebolavirus* and *Cuevavirus* genera indicated that L243/F245/L249 are identical or substituted conservatively for other hydrophobic amino acids that comprise part of the consensus NES sequence (L, V, I, F or M; Figure 4E); consistent with this, C-terminal NES activity was predicted for all of the proteins using NetNES (Figure S2C). In contrast, MABV (genus *Marburgvirus*) has a glutamine residue at the site corresponding to EBOV L243 (Figure 4E), and this results in a loss of predicted NES activity (Figure S2C). Thus, it appears that NES activity is important for species of the genus *Ebolavirus* and *Cuevavirus*, but not *Marburgvirus*. Interestingly, MABV VP24 is unique among the VP24 proteins of filoviruses in that it is reported not to bind Kαs (26, 44, 45). Thus, MABV VP24 would appear not to have the same requirement for interaction with nuclear trafficking machinery as the VP24 proteins of the other filoviruses.

### EBOV VP24 NES activity is not required for IFN/STAT1 antagonist function

The conservation of the NES sequence among ebolaviruses and LLOV indicates important function. Given the central role for VP24 in antagonising signalling by IFN/STAT1 (17-20), we assessed the effects thereon of NES mutations using an IFN-α/STAT1/2-dependent luciferase reporter gene assay, as previously (40, 46) (Figure 5). This indicated that GFP-VP24-NM potently inhibits IFN-α-induced luciferase expression, to an extent similar to that observed for WT VP24.

**Figure 5.**
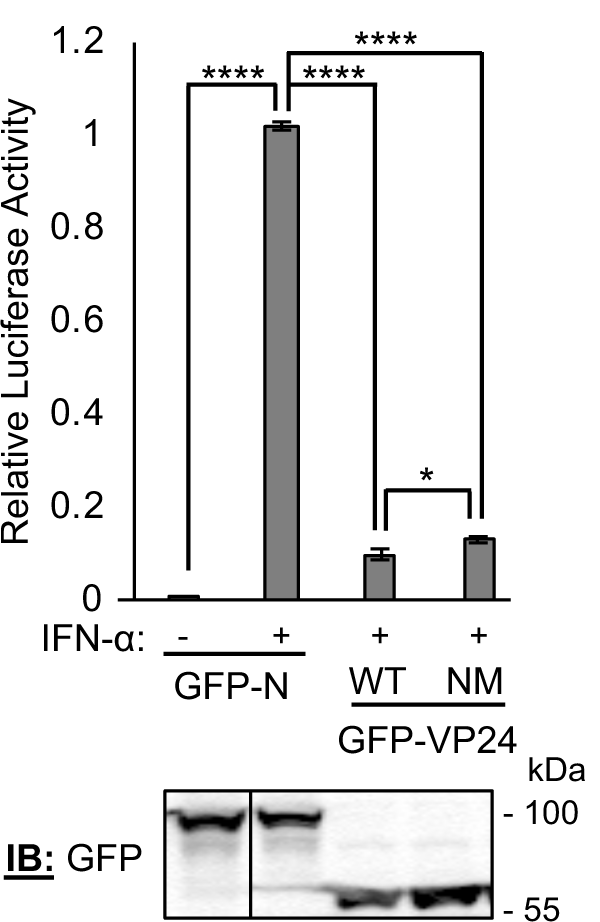
EBOV VP24 NES activity is not required for inhibition of IFN-α/STAT1/2-dependent gene expression. HEK293T cells co-transfected with pISRE-Luc and pRL-TK plasmids, and plasmids to express GFP-fused RABV N (control), VP24 WT or VP24 NM, were treated 8 h post-transfection with or without IFN-α (1,000 U/ml, 16 h) before determination of relative luciferase activity (mean ± SEM; n = 4 independent assays, each of which was performed in triplicate); *lower panel:* expression of proteins in cell lysates used in a representative assay was analysed by immunoblotting (IB) for GFP. Statistical analysis used Student’s *t*-test; *, p<0.05; ****, p < 0.0001.

To further examine effects of altered VP24 nuclear trafficking on STAT1 responses, we assessed nuclear import of STAT1 using CLSM analysis of COS7 cells expressing GFP-VP24 and immunostained for STAT1 following treatment without or with IFN-α and/or LMB. In agreement with results of the luciferase reporter assays, we observed that despite substantial re-localisation of GFP-VP24 to the nucleus in LMB-treated cells, IFN-α-dependent STAT1 nuclear localisation remained clearly inhibited (Figure S3). Together, these data indicate that nuclear export of VP24 is not required for inhibition of STAT1 responses, consistent with Kα binding representing the major antagonistic mechanism. Thus, it appears that active cytoplasmic re-localisation of VP24 principally enables other functions in cytoplasmic viral replication.

## Discussion

In this study we have shown that EBOV VP24 undergoes active trafficking between the nucleus and cytoplasm involving CRM1-dependent nuclear export *via* a NES at the VP24 C-terminus. The acquisition of active nuclear trafficking sequences is consistent with a requirement for highly regulated/dynamic localisation; furthermore, since VP24 is reported to oligomerise (potentially as tetramers) (38), it is likely that active nuclear trafficking is required for transport of VP24 multimers. The identified NES was not resolved in VP24 crystal structures (20, 47, 48) but localisation at the C-terminal end would be consistent with exposure and accessibility to CRM1 (20), and the predominantly cytoplasmic localisation of GFP-VP24 in resting cells suggests that the NES is the dominant trafficking signal at steady state. Intriguingly, previous studies indicated that a mutated VP24 protein defective for Kα-binding was more cytoplasmic than WT protein (49). This would be consistent with karyopherin binding mediating import; one might thus speculate that VP24 would require export mechanisms to enable cytoplasmic localisation/functions. Our findings are the first to confirm this is the case. Notably, the EBOV matrix protein VP40 has also been reported to localise to the nucleus in infected and transfected cells (16, 50); however, a direct role for active trafficking pathways to regulate localisation, distinct from mechanisms such as diffusion or interaction with other host factors, has not been defined. Thus, our data provides, to our knowledge, the first direct demonstration of a filovirus protein exploiting specific host trafficking machinery for nucleocytoplasmic transport, identifying a new mechanism in infection by these highly lethal pathogens.

Although the nucleus is not directly involved in the replication processes of most RNA viruses, proteins of a number of these viruses are reported to encode nuclear trafficking sequences, indicative of a requirement for dynamic regulation or specific accumulation in particular compartments. For example, the RABV IFN antagonist P protein encodes several NLSs and NESs (32, 43, 51-53), with regulatory mechanisms including co-localisation or overlap of the sequences, enabling co-regulation by mechanisms including phosphorylation (51-53). Although our data identify the C-terminal NES as a principal determinant of nucleocytoplasmic localisation of full-length VP24, the differential localisation and LMB sensitivity of VP24^1-88^ and VP24^89-172^, and the finding that VP24^89-251^ does not recapitulate nuclear accumulation of full-length VP24, suggest the presence of alternative regulatory sequences/mechanisms, potentially exposed by truncation. For example, VP24 is reported to associate with membranes (38), which might result in tethering within the cytoplasm under certain conditions. Interestingly, a recent study reported that sumoylation of residue K14 of VP24 enhances Kα binding and IFN antagonistic function (54). In contrast, ubiquitination, including at residue K206 within CL3 (Figure 2A), appears to negatively regulate IFN antagonist activity (54). Intriguingly, K14 is distal to CL1-3 but is within a predicted NES motif (Figure 2A). Whether NESs of VP24 undergo dynamic regulation by post-translational modification or other mechanisms will be of interest in defining the processes controlling immune evasion and replication by EBOV.

While some viral IFN antagonists use NESs to facilitate immune evasion, including through mislocalisation of associated STATs (33, 34), VP24 uses a mechanism of competitive binding to Kαs. Our finding that VP24 nuclear export is not required for STAT antagonism is consistent with this, and indicates that export relates to cytoplasmic roles including in nucleocapsid assembly and transport (4-14). The requirement for efficient translocation out of the nucleus is consistent with interaction of VP24 with Kα (see above), that underpins distinct functions in immune evasion. This is further supported by our finding that the C-terminal NES motif is conserved among VP24 of several filovirus species that have been shown to bind to Kαs or have conserved CL sequences (20, 26, 45), but not in MABV VP24 (Figure 4E, Figure S2C), which has no role in antagonising STAT1, and does not bind Kαs (26, 44).

Other than nucleocapsid formation and transport, cytoplasmic VP24 may also function in budding (5, 38, 55) as a ‘minor matrix protein’. Similarly to matrix proteins of a number of other viruses of the order *Mononegavirales*, VP24 is reported to have negative effects on transcription/genome replication (56, 57), likely due to roles in genome packaging/nucleocapsid condensation (7, 11, 56, 57). Interestingly, imaging of EBOV-infected cells indicated that VP24 accumulates within nucleoprotein-rich inclusion bodies only from 18 hours post-infection (16), presumably to permit sufficient transcription/replication before packaging for assembly and release. Thus, VP24 nucleocytoplasmic trafficking might provide regulation of the replication-assembly switch, similar to mechanisms proposed for dynamic nucleocytoplasmic localisation of matrix proteins of paramyxoviruses (31). Many matrix proteins of henipaviruses have also recently been shown to have specific intranuclear functions through interaction with nuclear/nucleolar proteins (36, 37). In light of our finding that VP24 undergoes nucleocytoplasmic trafficking, it is intriguing that mass spectrometry analysis identified a large number of proteins in the VP24 interactome with functions related to the nucleus (39), consistent with possible intranuclear functions of VP24. Given the nuclear localisation of EBOV VP40 early in infection (16), it appears that the nucleus might represent an important hub for EBOV-host interactions.

The multiple roles of EBOV VP24 are likely to account for the lack of success in generating a VP24-deficient virus (5); however, roles in virulence are indicated by the finding that mutations acquired in the VP24 gene during serial passaging in guinea pigs were necessary and sufficient to confer lethality (58). Notably, the adaptations were not associated with IFN antagonism (58), implying distinct roles in pathogenesis. Moreover, a phosphorodiamidate morpholino oligomer that targets VP24 mRNA protects rhesus monkeys against lethal EBOV infection (59, 60). Inhibition of CRM1-mediated nuclear export is reported to have antiviral effects against diverse viruses, including the RNA viruses Dengue virus and respiratory syncytial virus (61), while mutations impacting the RABV P protein NES correlate with attenuation *in vivo* (34). Thus, targeting VP24 regulatory mechanisms, including its nuclear export, may provide novel targets for anti-EBOV drug design.

## Materials and Methods

### Constructs, cells, transfections and drug treatments

The construct to express the minimal NLS from human cytomegalovirus UL44 protein (residues 425-433) fused to GFP was generated by subcloning from pEPI-GFP-UL44^425-433^ (42, 43) into the pEGFP-C1 vector C-terminal to GFP (Clontech). Constructs to express full-length or truncated EBOV-VP24 protein fused to GFP or GFP-UL44^NLS^ were generated by PCR amplification from pCAGGS-FLAG-VP24 (kindly provided by C. Basler, Georgia State University), and cloning into the pEGFP-C1 or pEGFP-C1-UL44^NLS^ vectors C-terminal to GFP/GFP-UL44^NLS^. NES mutations (L243A/F245A/L249A) were introduced into VP24 sequences by site-directed PCR mutagenesis, and cloning into vectors, as above. The construct to express FLAG-tagged Kα1 was a kind gift from C. Basler (Georgia State University). Other constructs have been described elsewhere (40).

COS7 and HEK293T cells were maintained in DMEM supplemented with 10 % FCS and GlutaMAX (Life Technologies), 5 % CO_2_, 37°C. Transfections used Lipofectamine 2000 and Lipofectamine 3000 (Invitrogen), according to the manufacturer’s instructions. To inhibit CRM1-dependent nuclear export pathway, cells were treated with 2.8 ng/ml LMB (Cell Signaling Technology and a gift from M. Yoshida, RIKEN, Japan) for 3 h. To activate STAT1 nuclear localisation, cells incubated in serum-free DMEM (with or without 2.8 ng/ml LMB, 3 h) were treated with IFN-α (Universal Type I IFN, PBL Assay Science; 1000 U/ml, 30 min).

### Confocal Laser Scanning Microscopy (CLSM)

For analysis of VP24 localisation (with or without Kα1 over-expression) or STAT1 localisation, cells growing on coverslips and treated with or without LMB and IFN-α were fixed using 3.7 % formaldehyde (10 min, room temperature (RT)) followed by 90 % methanol (5 min, RT) before immunostaining. Antibodies used were: anti-FLAG (Sigma-Aldrich, F1804), anti-STAT1 (Cell Signaling Technology, 14994), anti-mouse Alexa Fluor 568 (ThermoFisher Scientific, A11004), anti-rabbit Alexa Fluor 647 (ThermoFisher Scientific, A21244). For live cell imaging, cells growing on coverslips were analysed under phenol-free DMEM. Imaging used a Nikon C1 inverted confocal microscope with 63 X objective and heated chamber for live cells. Digitised confocal images were processed using Fiji software (NIH). To quantify nucleocytoplasmic localisation, the ratio of nuclear to cytoplasmic fluorescence, corrected for background fluorescence (Fn/c), was calculated for individual cells expressing transfected protein (40); the mean Fn/c was calculated for n ≥ 31 cells for each condition in each assay.

### Co-immunoprecipitation

Cells were transfected with plasmids and lysed for immunoprecipitation using GFP-Trap beads (Chromotek), according to the manufacturer’s instructions. Lysis and wash buffers were supplemented with cOmplete Protease Inhibitor Cocktail (Roche). Lysates and immunoprecipitates were analysed by SDS-PAGE and immunoblotting using antibodies against FLAG (Sigma-Aldrich, F1804), GFP (Roche Applied Science, 11814460001) and HRP-conjugated secondary antibodies (Merck). Visualisation of bands used Western Lightning chemiluminescence reagents (PerkinElmer).

### Luciferase Reporter Gene Assays

Cells were co-transfected with pISRE-Luc (in which Firefly luciferase expression is under the control of a STAT1/2-dependent *IFN-sensitive response element*-containing promoter) and pRL-TK (transfection control, from which *Renilla* luciferase is constitutively expressed), as previously described (46), together with protein expression constructs. Cells were treated 8 h post-transfection with or without IFN-α (1000 U/ml) before lysis 16 h later using Passive Lysis Buffer (Promega). Firefly and *Renilla* luciferase activity was then determined in a dual luciferase assay, as previously described (46). GFP-RABV N-protein, which does not affect STAT signalling, was used as a negative control, as previously (40). The ratio of Firefly to *Renilla* luciferase activity was determined for each condition, and then calculated relative to that for GFP-N-expressing cells treated with IFN-α (relative luciferase activity). Data from 4 independent assays were combined, where each assay result is the mean of three biological replicate samples.

### Statistical Analysis

Unpaired two-tailed Student’s *t*-test was performed using Prism software (version 7, GraphPad).

### Sequence analysis

VP24 protein sequences from *Zaire ebolavirus* (NCBI accession no. AGB56798.1), *Tai Forest ebolavirus* (YP_003815430.1), *Bundibugyo ebolavirus* (YP_003815439.1), *Sudan ebolavirus* (YP_138526.1), *Reston ebolavirus* (NP_690586.1), *Lloviu cuevavirus* (YP_004928142.1) and *Marburg marburgvirus* (ABE27080.1) were aligned using the COBALT constraint-based multiple alignment tool (NIH, NCBI). To identify potential NES sequences, VP24 protein sequences were analysed using the NetNES 1.1 server http://www.cbs.dtu.dk/services/NetNES/ (41).

## Data Availability

Data available upon request to Gregory W Moseley

## Acknowledgements

This research was supported by National Health and Medical Research Council Australia project grants 1160838, 1125704 and 1079211 (G.W.M), Australian Research Council discovery project grant DP150102569 (G.W.M) and Australian Government Research Training Program Scholarship (A.R.H). We acknowledge Cassandra David for assistance with tissue culture, and the facilities and technical assistance of the Monash Micro Imaging Facility (Monash University). Plasmids to express FLAG-VP24 and FLAG-Kα1 were kind gifts from C. Basler (Georgia State University). LMB was a gift from M. Yoshida (RIKEN, Japan).

