## Supplemental Figures for "The Ebola virus interferon antagonist VP24 undergoes active nucleocytoplasmic trafficking"

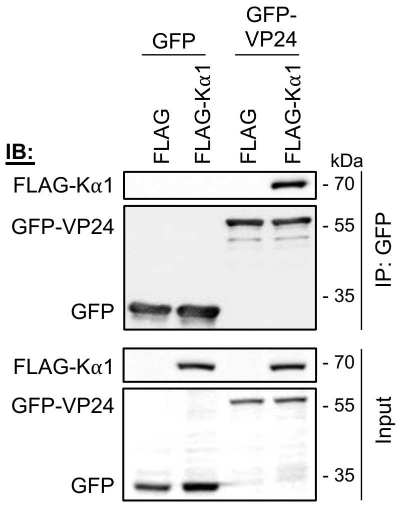


**Figure S1. Kα1 co-precipitates with EBOV VP24.** (A) HEK293T cells co-transfected to express GFP or GFP-VP24 with FLAG control or FLAG-Kα1 were lysed 24 h post-transfection before immunoprecipitation for GFP. Lysates (input) and immunoprecipitates (IP) were analysed by immunoblotting (IB) using antibodies against the indicated proteins. Results are representative of 3 independent assays.

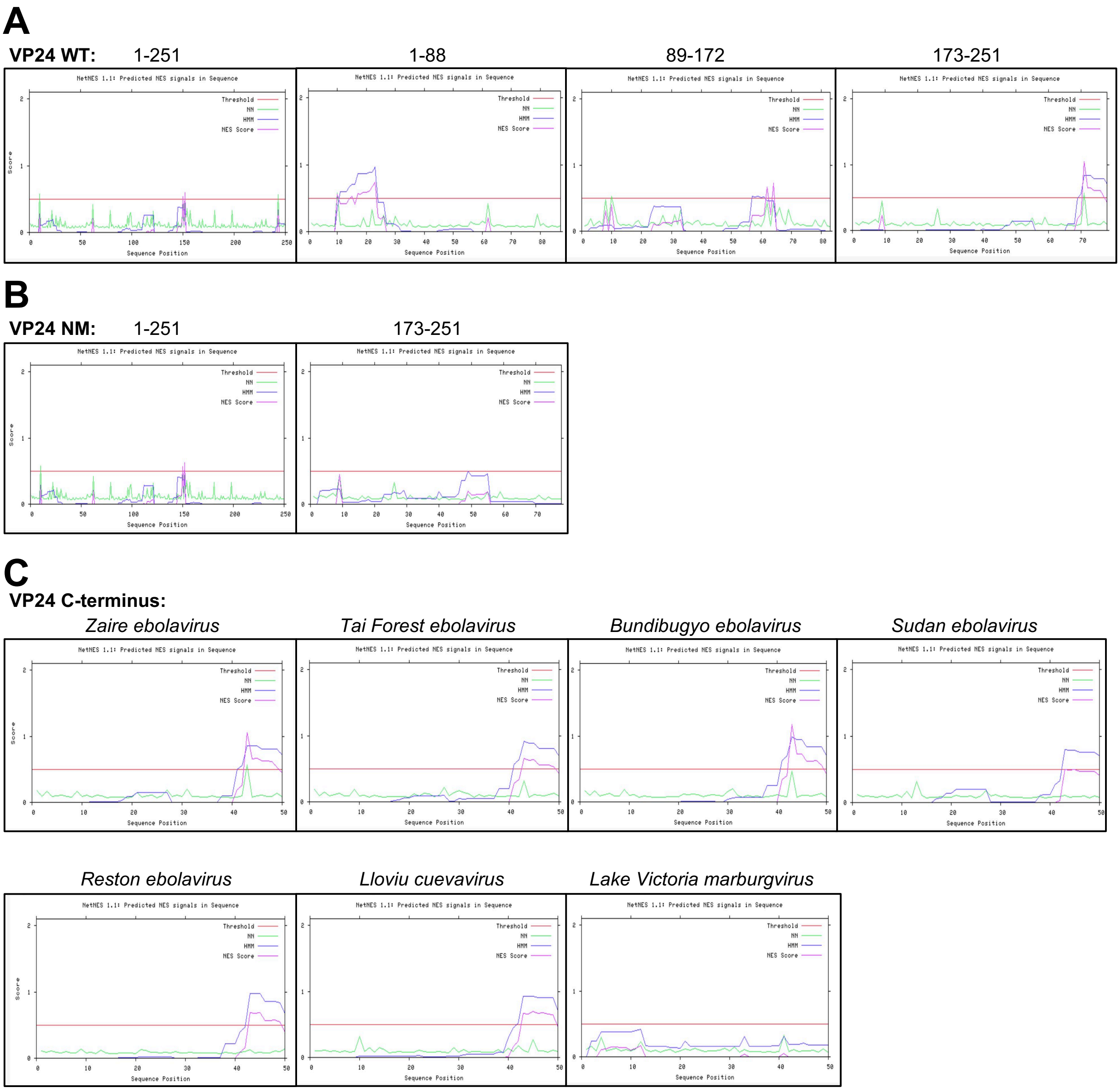


**Figure S2. Prediction of NES motifs in VP24 using NetNES.** (A) Analysis of full-length and truncated WT EBOV VP24 protein sequences using NetNES indicates four discrete sequences as potential NESs. (B) *In silico* substitution of residues L243, F245 and L249 for alanines (producing VP24 NM) results in loss of predicted NES function at the VP24 C-terminus. (C) NetNES analysis of C-terminal 51 residues of VP24 indicates conservation of the C-terminal NES among ebolaviruses and *Lloviu cuevavirus*, but not *Lake Victoria marburgvirus*. A NES is predicted if the NES score (pink line) exceeds the threshold (red line); the NES score is calculated based on Hidden Markov Model (HMM; blue line) and artificial Neural Network (NN; green line) scores (41).

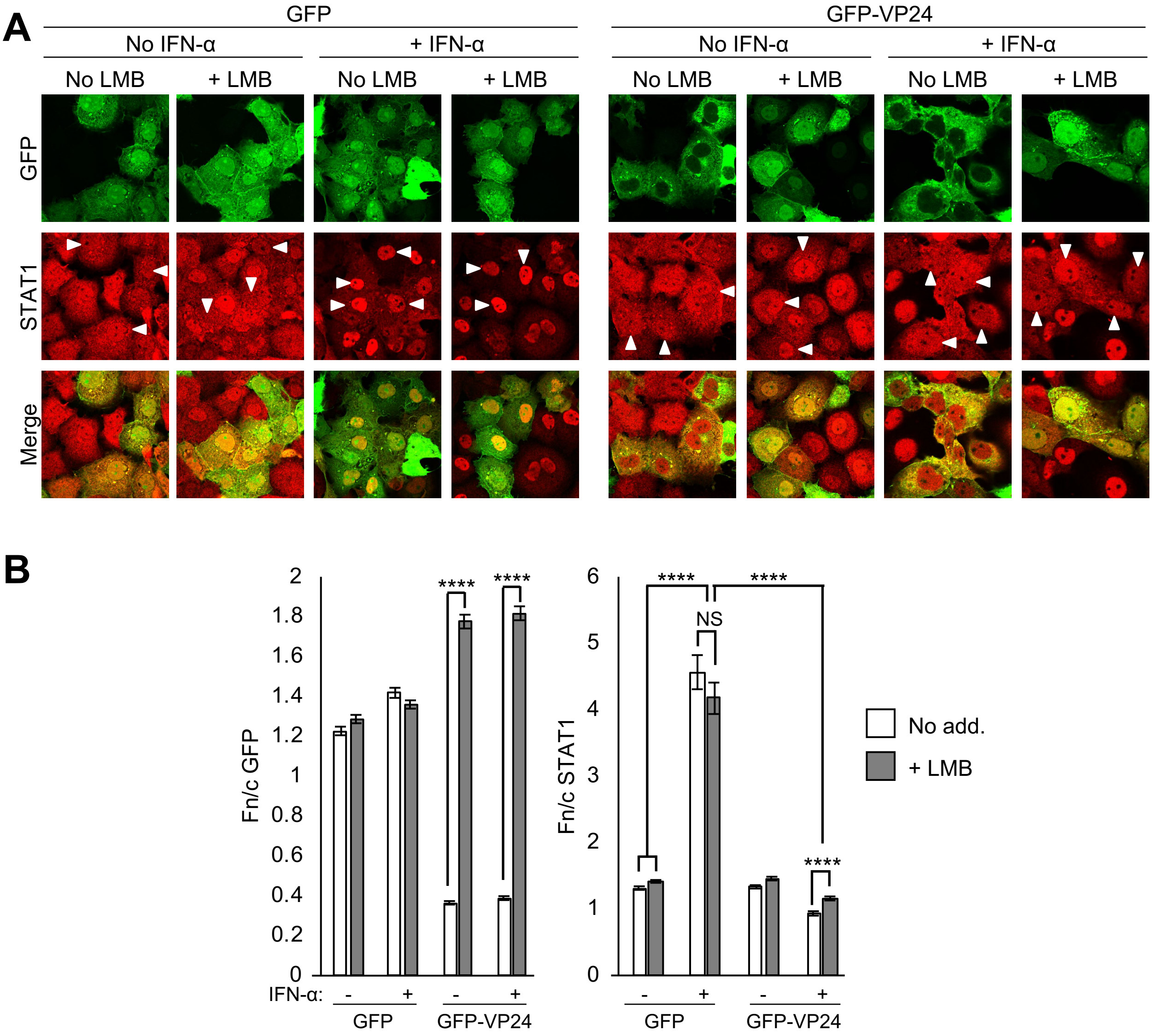


**Figure S3. Inhibition of nuclear export of VP24 does not prevent antagonism of STAT1.** (A) COS7 cells transfected to express GFP or GFP-VP24 were treated 24 h post-transfection with or without LMB (2.8 ng/ml, 3 h) and/or IFN-α (1000 U/ml, 30 min) before fixation, immunofluorescent staining for STAT1 (red) and CLSM analysis. Representative images are shown. Arrowheads indicate cells with detectable expression of GFP. (B) Images of transfected cells such as those shown in A were analysed to calculate the Fn/c for GFP and STAT1 (mean ± SEM, n ≥ 50 cells for each condition). Statistical analysis used Student's *t*-test; ****, $p < 0.0001$; NS, not significant; No add., no addition.
